## Supplementary Figures for "Tsetse salivary glycoproteins are modified with paucimannosidic *N*-glycans, are recognised by C-type lectins and bind to trypanosomes"

[1]Ludger Ltd., Culham Science Centre, Oxfordshire, OX14 3EB, UK. [2]Department of Vector Biology, Liverpool School of Tropical Medicine, Pembroke Place, Liverpool, L3 5QA, England, UK [3]Department of Tropical Disease Biology, Liverpool School of Tropical Medicine, Pembroke Place, Liverpool, L3 5QA, England, UK. [4]Laboratory of Microbiology, Parasitology and Hygiene (LMPH), University of Antwerp, Universiteitsplein 1, B-2610, Wilrijk, Belgium. [5]Department of Biomedical Sciences, Institute of Tropical Medicine Antwerp, Antwerp, Belgium

Running title: *Glycomics of tsetse fly saliva*

[#]These authors have contributed equally to this work and share first authorship

[□]Current address: Australian Research Council Centre of Excellence for Nanoscale Biophotonics, Macquarie University, Sydney, Australia

Keywords: glycosylation, mass spectrometry, tsetse fly, trypanosomiasis, insect saliva, hematophagy

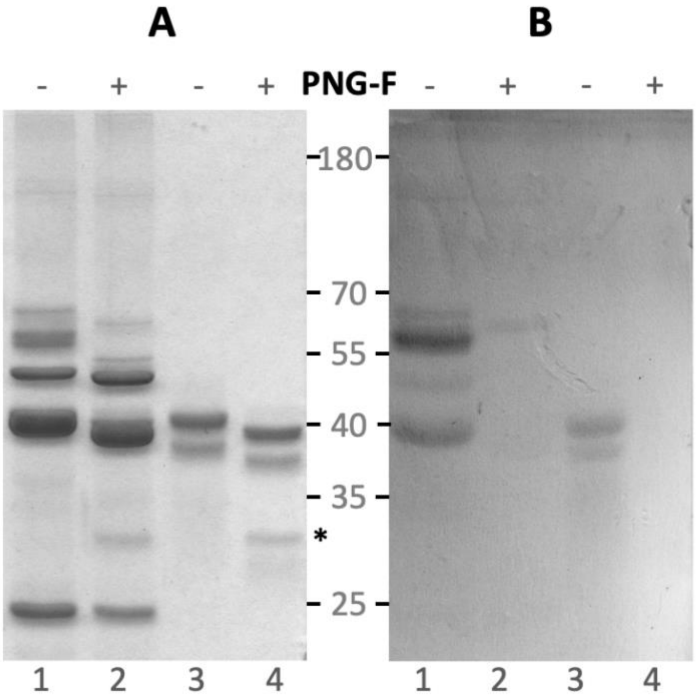


**Figure S1. Schiff's staining analysis of *G. morsitans* salivary glycoproteins.** 10 µg *G. morsitans* salivary proteins (lanes 1 and 2) and egg albumin (lanes 3 and 4) were incubated overnight with (+) or without (-) PNGase F to cleave *N*-glycans. Samples were resolved on a 12 % SDS-PAGE gel and stained with Colloidal Coomassie Blue (A) or Schiff's (B) staining. Asterisk indicates PNGase F enzyme.

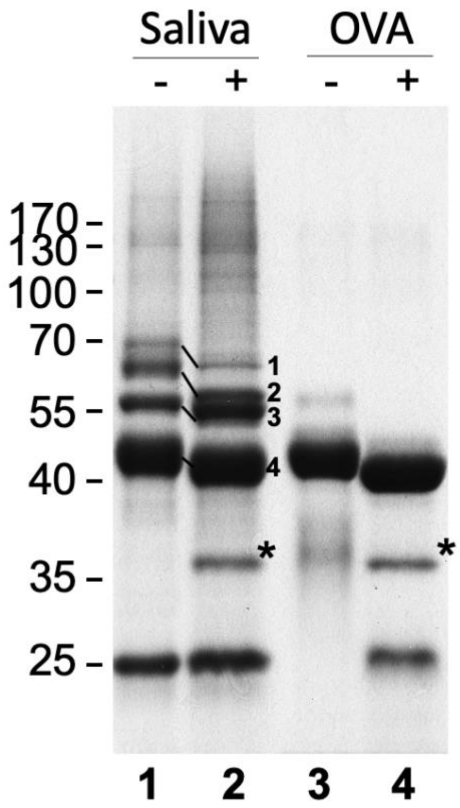


**Figure S2. Analysis of *G. morsitans* salivary glycoproteins.** 10 µg of *G. morsitans* salivary proteins (lanes 1 and 2) and 10 µg of egg albumin (lanes 3 and 4) were incubated overnight with (2 and 4) and without (1 and 3) PNGase F. After digestion, proteins were resolved by SDS-PAGE and Coomassie blue-stained. There was a notable shift in migration in 4 bands following PNGase F treatment. After in-gel trypsinization and MALDI-TOF MS analysis these bands were identified as 5' Nucleotidase (1), TSGF 2/Adenosine deaminase (2), TSGF 1 (3), and Tsal 1/2 (4). *, PNGase F enzyme.

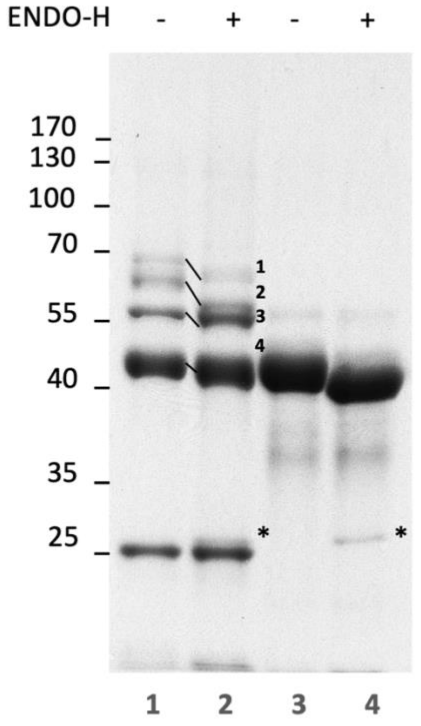


**Figure S3. Endo H cleavage of *G. morsitans* salivary glycoproteins.** 10 µg *G. morsitans* salivary proteins (lanes 1 and 2) and egg albumin (lanes 3 and 4) were incubated overnight with (+) or without (-) Endo H. Samples were resolved on a 12 % SDS-PAGE gel and Coomassie stained. There was a notable shift in migration in 4 bands (1-4) after deglycosylation. These bands were excised, trypsinised and identified by mass spectrometry. 1, 5' Nucleotidase; 2, TSGF 2/Adenosine deaminase; 3, TSGF 1; 4, Tsal 1/2. Asterisk indicates Endo H.

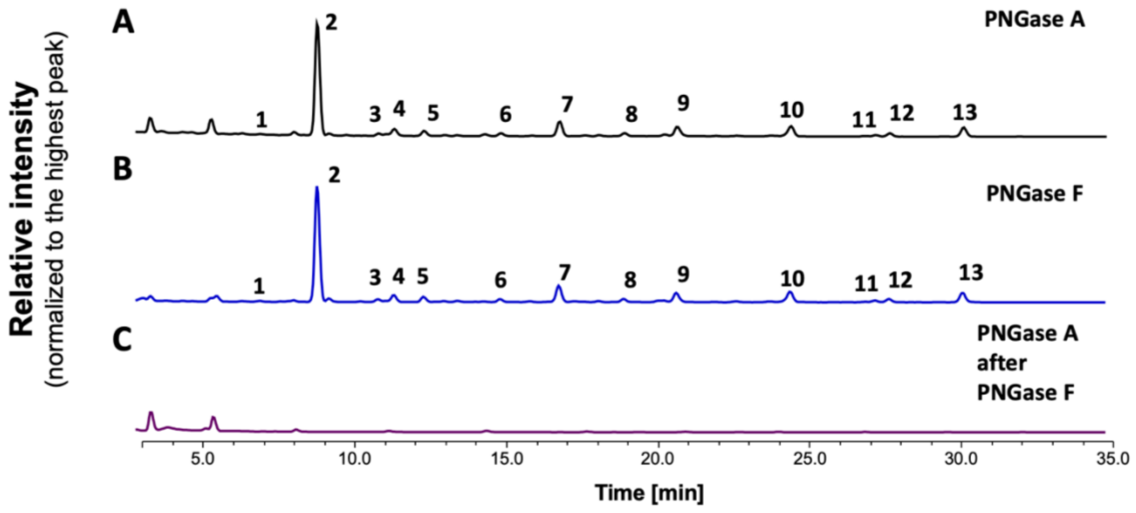


Figure S4. HILIC-(U)HPLC chromatograms of 2-AB labelled *N*-glycans released enzymatically from teneral fly saliva. (A) *N*-glycans released by PNGase A, (B) *N*-glycans released by PNGase F, (C) *N*-glycans released by PNGase A after deglycosylation with PNGase F. The glycan structures corresponding to the numbers on the HPLC peaks are listed in Table 3.

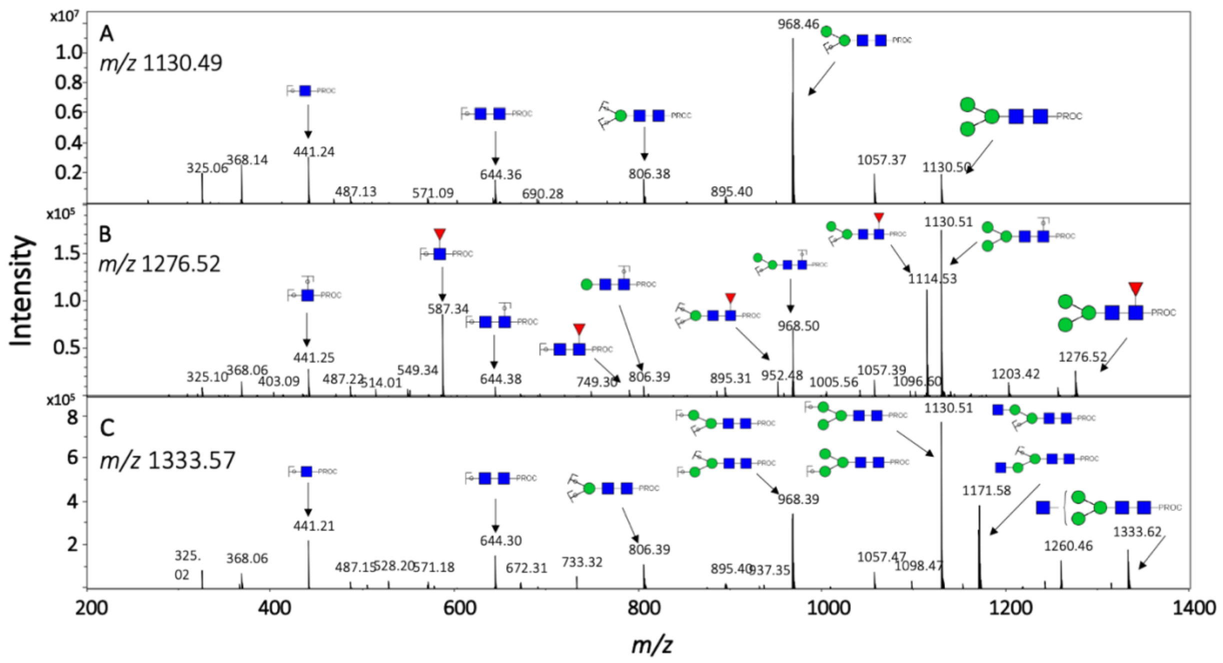


**Figure S5. Positive-ion ESI-MS/MS fragmentation spectra of procainamide-labelled *N*-glycans from teneral tsetse fly saliva.** Corresponding to (A) *m/z* 1130.49 ($Man_3GlcNAc_2$-Proc), (B) *m/z* 1276.52 ($Man_3GlcNAc_2Fuc$-Proc), (C) *m/z* 1333.57 ($Man_3GlcNAc_3$-Proc). Green circle, mannose; blue square, *N*-Acetylglucosamine; red triangle, fucose; Proc, procainamidxcxc

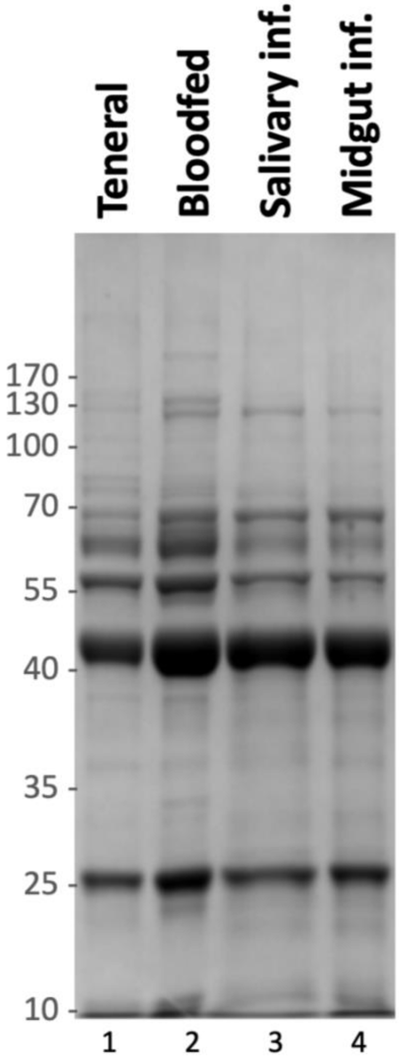


**Figure S6. SDS-PAGE fractionation of 10 µg of *G. morsitans* salivary profiles obtained from different infection stages**. Lane 1, teneral; Lane 2, 4-week old, bloodfed flies (Bloodfed); Lane 3, flies with salivary gland *T. brucei* infection (Salivary inf.); Lane 4, flies with midgut *T. brucei* infection (Midgut inf.).
