## Supplemental Table 3 for "Tsetse salivary glycoproteins are modified with paucimannosidic *N*-glycans, are recognised by C-type lectins and bind to trypanosomes"

### Supplementary Table 3. Proteomic identification of tsetse salivary proteins susceptible to Endo H treatment.

| Band number | Apparent molecular mass on SDS-PAGE | VectorBase Identifier | Name of protein in the band | Predicted molecular weight with signal peptide cleaved off | Predicted *N*-glycosylation sites (without signal peptide) | Unglycosylated sequons detected | % Peptide coverage |
|---|---|---|---|---|---|---|---|
| 1.2 | ≈65 kDa | GMOY012313-PA | 5’ Nuc[1] | ≈59 kDa | Asn85, Asn173, Asn270, Asn 440 | Asn173 | 4% |
| 2.2 | ≈58 kDa | GMOY012372-PA | TSGF2[2] | ≈56 kDa | Asn51, Asp103, Asn283, Asn347, Asp484 | Asn484 | 41% |
| | | GMOY012375-PA | Adenosine deaminase-related growth factor C[3] | ≈54 kDa | Asn19, Asn120, Asn171, Asn370, Asn454, Asn475, Asn483 | Asn120, Asn171, Asn454, Asn475 | 45% |
| 3.2 | ≈55 kDa | GMOY012373-PA | TSGF1[1] | ≈54 kDa | Asn339 | - | 61% |
| 4.2 | ≈42 kDa | GMOY012071-PA | Tsal1[4] | ≈44 kDa | Asn346 | - | 77% |
| | | GMOY012360-PA | Tsal2 (form B)[4] | ≈42 kDa | Asn238 | - | 63% |

[1] Caljon *et al*. (2010), [2] Li and Aksoy (2000), [3] Alves-Silva *et al.* (2010), [4] Li *et al*. (2000)
