## Supplemental Table 4 for "Tsetse salivary glycoproteins are modified with paucimannosidic *N*-glycans, are recognised by C-type lectins and bind to trypanosomes"

### Supplementary Table 4.

HILIC- LC-ESI-MS data with sugar composition and structures for N-glycans released by PNGase F and labelled with procainamide.
Hex, hexose; HexNAc, *N*-acetylhexoseamine
The symbols for glycan structures are adopted from the Consortium for Functional Glycomics:

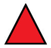 Fucose

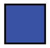 *N*-acetylglucosamine

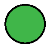 Mannose

| HPLC Peak Id | GU (Procainamide) | Structure | HILIC-LC-ESI-MS | | | | | | | |
|---|---|---|---|---|---|---|---|---|---|---|
| | | | Teneral Fly Saliva N-glycans Procainamide labelled | | | | | | | |
| | | | Composition | | | $[m/z]^{+}$ calculated | $[m/z]^{2+}$ calculated | $[m/z]^{+}$ registered | $[m/z]^{2+}$ registered | [m/z] characteristic fragment ions (composition) |
| | | | Hex | HexNAc | Fuc | | | | | |
| **1** | **3.21** | 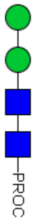 | **2** | **2** | **0** | 968.46 | 484.73 | 968.47 | nd | 441.20 (N-PROC)<br>644.33 (N2-PROC)<br>806.36 (H1N2-PROC) |
| **2** | **4.17** | 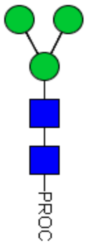 | **3** | **2** | **0** | 1130.51 | 565.76 | 1130.49 | 565.74 | 441.24 (N-PROC)<br>644.36 (N2-PROC)<br>806.38 (H1N2-PROC) |

| | | | | | | | | | | |
|---|---|---|---|---|---|---|---|---|---|---|
| | | | | | | | | | | 968.46 (H2N2-PROC) |
| **3** | **4.62** | 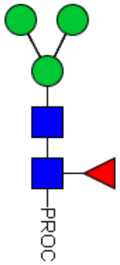 | **3** | **2** | **1** | 1276.57 | 638.79 | 1276.52 | 638.77 | 441.25 (N-PROC) 806.39 (H1N2-PROC) 1130.51 (H3N2-PROC)<br>587.34 (N1F1-PROC) 952.48 (H1N2F-PROC)<br>644.38 (N2-PROC) 968.50 (H2N2-PROC)<br>790.35 (N2F-PROC) 1114.53 (H2N2F-PROC) |
| **4** | **4.76** | 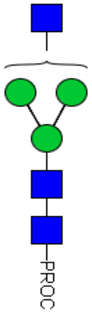 | **3** | **3** | **0** | 1333.59 | 667.30 | 1333.57 | 667.29 | 441.21 (N-PROC) 1130.51 (H3N2-PROC)<br>644.30 (N2-PROC) 1171.58 (H2N3-PROC)<br>806.39 (H1N2-PROC)<br>968.39 (H2N2-PROC) |
| **5** | **5.00** | 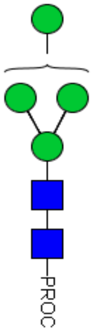 | **4** | **2** | **0** | 1292.56 | 646.78 | 1292.54 | 646.74 | 441.24 (N-PROC) 1130.50 (H3N2-PROC)<br>644.32 (N2-PROC)<br>806.38 (H1N2-PROC)<br>968.39 (H2N2-PROC) |
| **6** | **5.54** | | **4** | **3** | **0** | 1495.64 | 748.32 | 1495.65 | 748.30 | 441.24 (N-PROC) 1130.58 (H3N2-PROC) |

| | | | | | | | | | | |
|---|---|---|---|---|---|---|---|---|---|---|
| | | 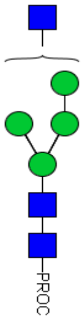 | | | | | | | | 644.36 (N2-PROC)<br>806.41 (H1N2-PROC)<br>968.51 (H2N2-PROC)<br>1171.57 (H2N3-PROC)<br>1292.61 (H4N2-PROC)<br>1333.62 (H3N4-PROC) |
| **7** | **6.00** | 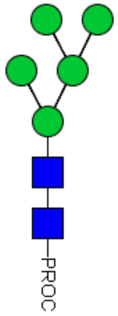 | **5** | **2** | **0** | 1454.61 | 727.81 | 1454.57 | 727.79 | 441.25 (N-PROC)<br>644.37 (N2-PROC)<br>806.38 (H1N2-PROC)<br>968.44 (H2N2-PROC)<br>1130.56 (H3N2-PROC)<br>1292.62 (H4N2-PROC) |

| | | | | | | | | | | |
|---|---|---|---|---|---|---|---|---|---|---|
| **8** | **6.46** | 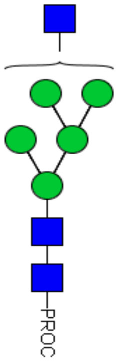 | **5** | **3** | **0** | 1657.69 | 829.35 | 1657.64 | 829.33 | 441.29 (N-PROC)<br>644.38 (N2-PROC)<br>806.40 (H1N2-PROC)<br>968.48 (H2N2-PROC)<br>1130.55 (H3N2-PROC)<br>1171.61 (H2N3-PROC)<br>1292.61 (H4N2-PROC)<br>1333.60 (H3N3-PROC)<br>1454.66 (H5N2-PROC)<br>1495.73 (H4N3-PROC) |

| | | | | | | | | | | |
|---|---|---|---|---|---|---|---|---|---|---|
| **9** | **6.87** | 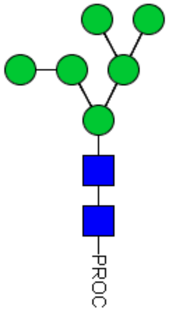 | **6** | **2** | **0** | 1616.67 | 808.84 | 1616.61 | 808.81 | 441.26 (N-PROC)<br>644.36 (N2-PROC)<br>806.39 (H1N2-PROC)<br>968.49 (H2N2-PROC)<br>1130.58 (H3N2-PROC)<br>1292.62 (H4N2-PROC)<br>1454.70 (H5N2-PROC) |
| **10** | **7.79** | 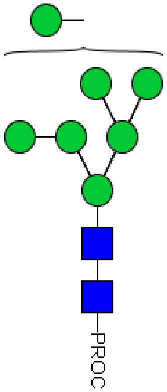 | **7** | **2** | **0** | 1778.72 | 889.86 | 1778.68 | 889.84 | 441.25 (N-PROC)<br>644.36 (N2-PROC)<br>806.46 (H1N2-PROC)<br>968.55 (H2N2-PROC)<br>1130.55 (H3N2-PROC)<br>1292.62 (H4N2-PROC)<br>1454.70 (H5N2-PROC)<br>1616.75 (H6N2-PROC) |
| **11** | **8.53** | 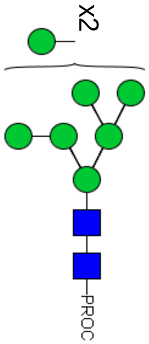 | **8** | **2** | **0** | 1940.77 | 970.89 | nd | 970.87 | 441.23 (N-PROC)<br>644.29 (N2-PROC)<br>806.49 (H1N2-PROC)<br>852.27 (H1N4)<br>1014.35 (H1N6)<br>1130.45 (H3N2-PROC)<br>1176.41 (H1N5)<br>1292.62 (H4N2-PROC)<br>1500.54 (H1N8) |

| | | | | | | | | | | |
|---|---|---|---|---|---|---|---|---|---|---|
| | | | | | | | | | | 968.54 (H2N2-PROC) 1338.42 (H1N7) |
| **12** | **8.66** | 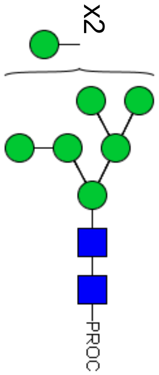 | **8** | **2** | **0** | 1940.77 | 970.89 | nd | 970.87 | 441.24 (N-PROC) 1014.30 (H1N6) 1454.59 (H5N2-PROC)<br>644.33 (N2-PROC) 1130.58 (H3N2-PROC) 1500.44 (H1N8)<br>806.38 (H1N2-PROC) 1176.42 (H1N5)<br>852.31 (H1N4) 1292.50 (H4N2-PROC)<br>968.44 (H2N2-PROC) 1338.42 (H1N7) |
| **13** | **9.35** | 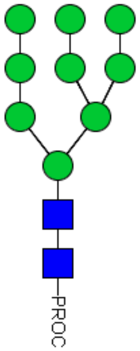 | **9** | **2** | **0** | 2102.83 | 1051.92 | nd | 1051.90 | 441.28 (N-PROC) 1014.08 (H1N6) 1454.51 (H5N2-PROC)<br>644.35 (N2-PROC) 1130.51 (H3N2-PROC) 1500.49 (H1N8)<br>806.39 (H1N2-PROC) 1176.40 (H1N5) 1662.53 (H1N9)<br>852.25 (H1N4) 1292.53 (H4N2-PROC)<br>970.86 (H2N2-PROC) 1338.38 (H1N7) |
