## Supplemental Table 5 for "Tsetse salivary glycoproteins are modified with paucimannosidic *N*-glycans, are recognised by C-type lectins and bind to trypanosomes"

Supplementary Table 5

HILIC-LC-ESI-MS/MS data for N-glycans released by PNGaseF. Table shows details for three representative structures from Teneral Fly Saliva.

* Mass corresponds to o loss of diethylamine ion (73 Da) [1].

| HILIC Peak ID | Structure | Composition | | | $[m/z]^+$ calculated | $[m/z]^+$ registered | [m/z] characteristic fragment ions (composition) |
|---|---|---|---|---|---|---|---|
| | | Hex | HexNAc | Fuc | | | |
| 2 | 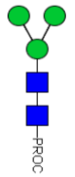 | 3 | 2 | 0 | 1130.51 | 1130.49 | 325.06 (H2) 571.09 (N2-PROC)* 806.38 (H1N2-PROC)<br>368.14 (N-PROC)* 644.36 (N2-PROC) 895.40 (H2N2-PROC)*<br>441.24 (N-PROC) 733.14 (H1N1-PROC)* 968.46 (H2N2-PROC) |
| 3 | 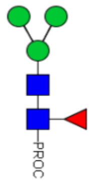 | 3 | 2 | 1 | 1276.57 | 1276.52 | 325.10(H2) 571.22 (N2-PROC)* 790.35 (N2F-PROC) 968.50 (H2N2-PROC) 1130.51 (H3N2-PROC)<br>368.06 (N-PROC)* 587.34 (N1F1-PROC) 806.39 (H1N2-PROC) 1041.41 (H2N2F-PROC)*<br>441.25 (N-PROC) 644.38 (N2-PROC) 895.31 (H2N2-PROC)* 1057.39 (H3N2-PROC)*<br>514.00 (N1F1-PROC)* 733.09 (H1N2-PROC)* 952.48 (H1N2F-PROC) 1114.53 (H2N2F-PROC) |
| 4 | 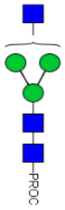 | 3 | 3 | 0 | 1333.59 | 1333.57 | 325.02 (H2) 644.30 (N2-PROC) 968.39 (H2N2-PROC) 1260.46 (H3N3-PROC)*<br>368.06 (N-PROC)* 733.32 (H1N2-PROC)* 1057.47 (H3N2-PROC)* 1171.58 (H2N3-PROC)<br>441.21 (N-PROC) 806.39 (H1N2-PROC) 1098.47 (H2N3-PROC)*<br>571.18 (N2-PROC)* 895.40 (H2N2-PROC)* 1130.51 (H3N2-PROC) |

1 Kozak, R. P., Tortosa, C. B., Fernandes, D. L. and Spencer, D. I. (2015) Comparison of procainamide and 2-aminobenzamide labeling for profiling and identification of glycans by liquid chromatography with fluorescence detection coupled to electrospray ionization-mass spectrometry. Anal Biochem. **486**, 38-40
